# PIGOME: An Integrated and Comprehensive Multi-omics Database for Pig Functional Genomics Studies

**DOI:** 10.1101/2024.03.10.583139

**Authors:** Guohao Han, Peng Yang, Yongjin Zhang, Qiaowei Li, Xinhao Fan, Ruipu Chen, Chao Yan, Mu Zeng, Yalan Yang, Zhonglin Tang

## Abstract

In addition to being a major source of animal protein, pigs are important model for the study of development and diseases in humans. During the past two decades, thousands of high-throughput sequencing studies in pigs have been performed using a variety of tissues from different breeds and developmental stages. However, the multi-omics database specifically used for pig functional genomic research is still limited. Here, we present a user-friendly database of pig multi-omics named PIGOME. PIGOME contains seven types of pig omics datasets, including whole-genome sequencing, RNA-seq, miRNA-seq, ChIP-seq, ATAC-seq, bisulfite-seq, and MeRIP-seq, from 6,901 samples and 392 projects with manually curated metadata, integrated gene annotation, and quantitative trait locus information. Furthermore, various ‘explore and browse’ functions have been established for user-friendly access to omics information. PIGOME implemented several tools to visualize genomic variants, gene expression, and epigenetic signals of a given gene in the pig genome, enabling efficient exploration of spatial-temporal expression/epigenetic pattern, function, regulatory mechanism, and associated economic traits. Collectively, PIGOME provides valuable resources for pig breeding and is helpful for human biomedical research. PIGOME is available at https://pigome.com.

## Introduction

Pig production accounts for a large proportion of the animal husbandry economy and is one of the mainstays of the global agricultural economy [1,2]. Moreover, pigs have been shown to be an important biomedical model for the study on human development and diseases [3–5]. Local adaptation and artificial selection have resulted in significant phenotypic differences and genetic diversity in pigs [6]. It is an exceptional model to elucidate the underlying mechanisms of traits, such as meat production, litter size, coat color, immune and diseases [7,8]. In the past several decades, with the development of advanced sequencing technologies, massive amounts of high-throughput sequencing data have been generated at multi-omics layers. These massive data provide a better understanding of evolution, selection, trait formation, development, and diseases in the genetic mechanisms and uncover many key variants, genes, and regulatory elements regulating various biological processes and associated with economic traits in pigs [9–11]. Our recent study based on high-resolution DNA methylome and transcriptome in skeletal muscle at 27 developmental stages provided insights into the molecular regulation of skeletal muscle development and diversity and uncovered candidate genes, such as *IGF2BP3* and *SATB2*, which contributes to skeletal muscle development, that provides a representative case to integrate multi-omics data facilitating the functional genomics studies of pigs [6,12]. Therefore, it is necessary to integrate multi-omics data to support the scientific discovery of pig genetics and breeding.

In particular, a mounting number of high-throughput sequencing studies in pigs have been performed based on a variety of tissues from different breeds and developmental stages [13–17]. However, these datasets are generated from different laboratories and sequencing platforms, making their retrieval, management, standard processing, and visualization time-consuming and difficult [18]. Furthermore, mining and integrated analysis of these datasets to explore the biological functions and regulatory mechanism remains a challenge [19]. Over the past several years, a limited number of specialized pig-related databases have been developed. Recently, IAnimal (https://ianimal.pro/) [20] was released, which includes pig multi-omic and genome annotation information. Similarly, ISwine (http://iswine.iomics.pro/) [21] contains published pig genome, transcriptome, and QTX data, similar to quantitative trait locus (QTL) information. To date, the Animal Omics Database (http://animal.nwsuaf.edu.cn/) contains only pan-genome sequencing datasets. However, the analysis and visualization abilities of these databases are limited (see Table 2). There is still a lack of comprehensive multi-omics databases dedicated to functional genomic research on pigs.

**Table 1.** Summary of omics data in PIGOME database.

| Type | Sample | Project | Tissue | Breed | Development<br>al stage | Data amount | Characteristic |
| --- | --- | --- | --- | --- | --- | --- | --- |
| RNA-seq | 4217 | 268 | 50 | 74 | 29 | 33.92 Tb | 31,908 genes |
| miRNA-seq | 995 | 78 | 39 | 32 | 29 | 269.35 Gb | 544 miRNAs |
| ATAC-seq | 58 | 5 | 13 | 6 | 5 | 1.26 Tb | 2,884,709 peaks |
| Bisulfite-seq | 309 | 20 | 26 | 13 | 9 | 2.17 Tb | 34,560,764 CpG sites |
| ChIP-seq | 388 | 16 | 22 | 6 | 7 | 2.75 Tb | 21,318,546 peaks |
| MeRIP-seq | 47 | 5 | 5 | 5 | 10 | 174.32 Gb | 424,376 peaks |
| WGS | 887 | - | - | 53 | - | 8.67 Tb | 12,074,987 SNPs |

*Note:* WGS, whole genome sequencing.

**Table 2.** Comparison of PIGOME with other databases.

|  | PIGOME | IAnimal-pig | Iswine | AOD |
| --- | --- | --- | --- | --- |
| Data type | WGS, RNA-seq, miRNA-seq, ChIP-seq, ATAC-seq, BS-seq MeRIP-seq | WGS, RNA-seq, ATAC-seq, ChIP-seq, | WGS, RNA-seq | WGS |
| Sample | 6901 | 10714 | 4107 | 12 |
| Data volume (TB) | 49.21 | 132.92 | Not provide | Not provide |
| Breed | 113 | Statistical difficulties | ~23 | 12 |
| Tissue | 71 | ~130 | 95 | Not provide |
| Developmental stage | 29 | Statistical difficulties | ~80 | Not provide |
| Analysis tools | Jbrowse2, IGV, Get Sequence, Primer Design, BLAST, Gene Network, Target Prediction, Find Tissue-specific Genes, API | Jbrowse, BLAST, Primer, Gene Network, Gene Correlation Coefficient, Signal Plotter, Signal Comparison, Genotype Plotter, Enrichment, API | Jbrowse, Primer, BLAST, Prioritize | GBrowse, BLAST, BLAT |

*Note:* WGS, whole genome sequencing. Data collected until December 2023.

To address these challenges, we developed PIGOME, an integrated and comprehensive web database containing seven types of sequencing data from 6,901 datasets and 392 projects, which is currently the most comprehensive omics database for pigs. PIGOME allows researchers to explore and utilize pig multi-omics data easily and effectively. Specifically, PIGOME supports the exploration, analysis, and visualization of genomic variations, expression patterns, regulatory networks, and epigenetic modifications of annotated and predicted pig genes (protein-coding genes (PCGs), microRNAs (miRNAs), and circular RNAs (circRNAs)). PIGOME also supports a tissue-specific analysis tool that allows users to identify the characteristics of genes, miRNAs, and circRNAs in specific tissues. In addition, PIGOME deploys nine tools, such as JBrowse [22], IGV [23], and Sequenceserver [24] to enable users to upload their files to visualize epigenetic signals and perform sequence alignment across the genomes of different pig breeds. In summary, PIGOME is an important resource for pig functional genomics studies and will be of interest to a broad readership in the fields of animal genetics, breeding, and biomedical research.

## Data collection and database construction

### Data collection

Seven types of high-throughput sequencing data (whole-genome sequencing [WGS], transcriptome sequencing [RNA-seq], microRNA sequencing [miRNA-seq], chromatin immunoprecipitation sequencing [ChIP-seq], assay for transposase-accessible chromatin sequencing [ATAC-seq], bisulfite sequencing for DNA methylation analysis [BS-seq] and methylated RNA immunoprecipitation sequencing [MeRIP-seq]) of pigs were collected from NCBI Sequence Read Archive (SRA, https://www.ncbi.nlm.nih.gov/sra/) and CNCB Genome Sequence Archive (GSA, https://ngdc.cncb.ac.cn/gsa/). BS-seq contains two types of data, including whole genome bisulfite sequencing (WGBS) and reduced representation bisulfite sequencing (RRBS).

WGS datasets were employed to construct the pan-genome and identify genomic variants, including single nucleotide polymorphisms (SNPs) and DNA insertion and deletion (InDels), within the genome. RNA-seq and miRNA-seq datasets were used to analyze the expression of mRNA and ncRNAs (such as miRNA, lncRNA, and circRNAs). ChIP-seq datasets were used to identify the modification of CTCF, histone modifications, and POL2 across the genome. The ATAC-seq datasets were used to identify open chromatin regions across the genome. BS-seq and MeRIP-seq datasets were used to analyze genome-wide DNA and RNA methylation, respectively.

All these datasets were manually collected with all the relevant metadata for fast and accurate data retrieval and statistical analysis, including project ID, tissue, developmental stage, breed, read number, platform, references, and others of the same kind. For the developmental stage information, referring to our previous paper [12], it was classify into 27 known stages (E33 to D180) and two fuzzy developmental stages ‘Unknown’ and ‘Adult’. Samples with poor data quality (mapping rate < 30% and data volume < 0.15 Gb) were excluded manually. To explore gene functions more conveniently, gene annotation information was integrated from Ensembl 100 [25], miRbase 22.1 [26] and eggNOG 5 [27], while QTL information was integrated from Animal QTLdb 46 [28] (Figure 1).

**Figure 1.**
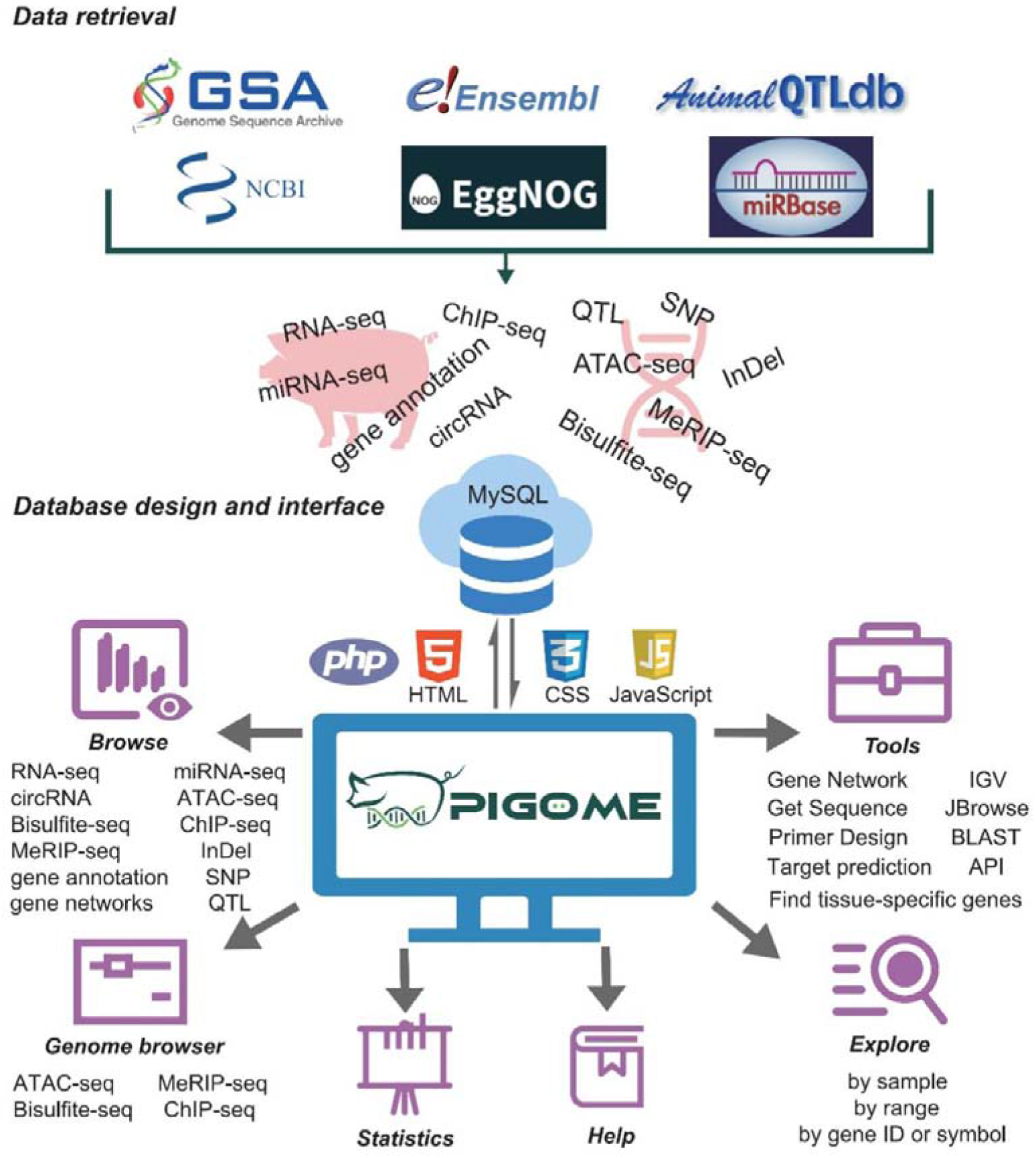
Database contents and construction. The present version of PIGOME contains 7 types of omics data, gene annotation and QTL information in pigs. PIGOME also contains practical functions and analytical tools to browse, explore and visualize omics data. QTL, quantitative trait locus.

### Data processing

The FASTA file of *Sus scrofa* reference genome (build 11.1) and the GTF annotation file (release 100) were downloaded from the Ensembl database. All raw FASTQ files were downloaded from the NCBI SRA and CNGB GSA databases. Fastp (v0.20.0) [29] was used to trim and filter the raw reads. For RNA-seq, HISAT2 (v2.0.5) [30] was used to map to the reference genome. Gene expression quantification in transcripts per kb exon per million mapped reads (TPM) was calculated using StringTie (v1.3.6) [31]. For miRNA-seq, adapters were removed using Cutadapt (v1.8. dev0) [32]. After remove adapters, these reads were then aligned to annotated pig miRNAs gathered from miRBase [26] and the pig reference genome using miRDeep2 (v0.1.2) [33]. For circRNAs, HISAT2 (v2.0.5) was used for alignment with the reference genome. Novel circRNAs were identified using CIRIquant (v1.1.1) [34] and Find_circ (v2) [35]. For ATAC-seq and ChIP-seq, all reads were mapped using Bowtie2 (v2.3.5.1) [36]. The peaks were identified using MACS2 (v2.2.6) and annotated using SnpEff (v4.2) [37]. Bismark (v0.23.0) [38] was used to align the reference genome using default parameters. All CpG sites were identified and annotated as previously described [12]. For MeRIP-seq, all reads were mapped using HISAT2 (v2.0.5), and peaks were identified using exomePeak2 (v2) [39] and annotated using SnpEff (v4.2). Picard (v2.25.7) was used to remove duplicate PCR reads for ATAC-seq, ChIP-seq, and MeRIP-seq. BigWig files for genome browser visualization were generated using deepTools (v3.5.1) [40] and bedGraphToBigWig (v4). For WGS, all reads were aligned to the reference genome using BWA (v0.7.12) [41]. The SNP and InDel calling used a Unified Genotyper approach as implemented in the GATK package (v4.1.5.0). Considering that the volume of SNP data is too large, SNPs in intergenic regions was not shown in the website, but could be accessed from the corresponding authors upon reasonable request. Furthermore, tissue-specific genes were identified using the R package TissueEnrich (v3.15) [42] based on the expression matrix of mRNA and ncRNAs. The rcorr function of Hmisc (v5.1-1) in R package was used to calculate the expression correlation between mRNA, miRNA and circRNA to construct the co-expression networks with r > 0.85 and P value < 0.01 as thresholds. The intersection results of RNAhybrid (v2.1.2) [43] and miRanda (v3.3a) [44] were used to predict putative targets for mRNAs and circRNAs with E value < -20.

### Website implementation

PIGOME was built by Thinkphp 6.0.12 (https://www.thinkphp.cn/), a mature model-view-controller (MVC) framework, deployed in CentOS 7.9 system. All omics data were stored in MySQL 5.6.50 (https://www.mysql.com/). Web interfaces were developed using HTML, CSS, JavaScript and Bootstrap 5.0.2 (https://getbootstrap.com/). Most of the interactive charts and tables were implemented with ECharts 5.3.1 (https://echarts.apache.org/) and Bootstrap Table 1.14.2 (https://bootstrap-table.com/) (Figure 1). Network proxy services were provided through Nginx 1.20.1 (https://www.nginx.com/). We recommend visiting PIGOME using Google Chrome, Microsoft Edge, or Mozilla Firefox.

## Database content and usage

### Data collection and statistics

At present, PIGOME v1.0 collects 7 types of multi-omics datasets in pigs, including WGS, RNA-seq, miRNA-seq, ChIP-seq, ATAC-seq, BS-seq (WGBS and RRBS) and MeRIP-seq. It contained 6,901 samples from 392 projects, including 113 breeds, 71 tissues, and 29 developmental stages. The total clean data reached 49.21 Tb (Table 1 and 2). The RNA-seq database represents the most abundant datasets in our database, including 4,217 samples, 74 breeds, 50 tissues, and 29 developmental stages (Table 1). To better interpret the omics data, we integrated 32,452 gene annotations and 29,687 QTLs. Gene annotation information records commonly have 22 attributes, including gene symbol, gene type, description, muscle biology, GO, KEGG, CAZy, and PFAM. In addition, the QTL information collected 11 attributes, mainly including position, QTL ID, name, type, trait, and PubMed ID. Additional statistics are summarized on the statistics page (https://pigome.com/statistics.html).

### PIGOME features and functions

PIGOME includes genomics (SNPs, InDels, and genome annotation), epigenomics (chromatin accessibility, histone modifications, DNA/RNA methylation), and transcriptomics (the abundance of mRNA and ncRNAs). In addition, it provides useful and user-friendly functions to help users perform advanced analyses (Figure 1).

#### Browse

Users can easily browse omics data using a Browse tag in the toolbar. After clicking on the omics data type, a summary information related to the data will be displayed. On the summary page of each level of omics data, users can obtain specific statistical data, including sample, gene, and other related information, and freely download these tables and charts. For more details, users can click the icon in the ‘Details’ column of a given gene or sample information table on the page, which links to the gene expression page. Basic information about the gene or sample is placed at the top of the gene expression page, with links to external databases. Different gene expression pages contained different sections. Specifically, the gene expression pages of RNA-seq, miRNA-seq, and circRNA can show the gene TPM values in various tissues, breeds, or developmental stages in given tissues, also display the TPM values of subgroups of samples freely selected by users. In addition, to better understand the gene function, it integrates a variety of gene annotations. For better finding the co-regulation between genes, the page shows the gene network of query gene. The expression pattern of a user given gene can be visualized in boxplots, bar charts, and line charts, moreover, the data will be conveniently presented in a table below the graph. Furthermore, the detail page of ATAC-seq, bisulfite-seq, ChIP-seq, and MeRIP-seq provides an IGV genome browser and a table to display information, allowing users to freely explore any genome intervals of each sample. Additionally, users can browse the allele frequency of a certain loci in different breeds on the detail page of SNP and InDel using a bar chart and table. This page also provides QTL information related to this region. In summary, PIGOME has a variety of browsing functions, paving the way for the integration and investigation of different omics in pigs.

#### Explore

For convenient usage, PIGOME provides three search engines to explore the whole database, including ‘by gene ID or symbol’, ‘by range’, and ‘by sample’. For ‘by gene ID or symbol’, users can explore by inputting gene ID or symbol. On the ‘by range’ page, users can explore by selecting chromosomes and entering the starting position and ending position. On the results page of ‘by gene ID or symbol’ and ‘by range’, all datasets related to a given gene or a given range are integrated and displayed. Additionally, users can obtain more detailed information by clicking on the links in the table. For ‘by sample’, users can fuzzily explore by selecting the dataset and entering SRR ID, Sample ID, or Project ID. Furthermore, the results page of ‘by sample’ displays the relevant sample information and provides relevant links. Importantly, all figures and data of the search results can be downloaded and edited easily.

#### Genome browser

PIGOME embeds a custom genome browser based on JBrowse2 to help users compare and analyze various omics datasets. PIGOME contains information on the sequence and gene annotations from Ensembl. By inputting the genome range or gene ID, users can explore the omics data related to the gene of interest. All tracks were marked according to the type of omics data, tissue, breed, and developmental stage. In the tracking group, the tracks of interest can be displayed by switching the checkboxes.

#### Tools

We have integrated nine practical tools, including IGV, JBrowse, ‘Get Sequence’, ‘Primer Design’, BLAST, ‘Gene Network’, ‘Target prediction’ ‘Find tissue-specific genes’, and API (Table 2). For IGV and JBrowse, users can check, verify, and interpret their own sequencing and genome data online. For ‘Get Sequence’, users can quickly extract the required gene sequence from large number of nucleotide sequences. Then, ‘Primer Design’ tools [45] can help users to design primers from DNA/RNA sequences of interest for further experimental verification. For BLAST, users can perform an alignment analysis based on their own sequences with 23 pig genomes. ‘Gene Network’ tool is helpful to find the gene regulatory network formed by the interaction between genes. For ‘Target prediction’, users can explore the regulatory role of miRNA in gene expression and find potential functional miRNA associated with economical traits. Furthermore, a tool called ‘Find tissue-specific genes’ was developed, that helps users quickly find tissue-specific genes, miRNAs, and circRNAs based on our massive expression data. Additionally, for API tools, users with basic programming skills can obtain the omics data more flexibly and explore functional genes more effectively. These tools will assist us to better explore the biological mechanisms of various biological processes and important economic traits in pigs.

Additionally, users can easily find more help from the database through the ‘Help tag’ in the toolbar.

### Comparison with the existing databases

To date, several user-friendly databases have been established to aggregate multi-omics datasets in pigs (Table 2). IAnimal [20] is a multi-species and multi-omics database, encompassing four types of pig omics data, including WGS, RNA-Seq, ChIP-Seq and ATAC-Seq. ISwine [21] serves as a professional pig omics database offering access to WGS, RNA-seq, quantitative traits and annotation information. While both IAnimal and ISwine provide valuable sample meta-information, including details on tissue, developmental stage, and breed, they lack secondary classification and correction of this metadata. Consequently, comparative analyses of gene expression regulation between developmental stages or breeds become challenging. The AOD database focuses on providing 12 *de novo* genome assemblies in pigs. However, PIGOME emerges as a standout platform in this landscape. Notably, it boasts the widest array of omics data types and meticulously curated sample meta-information. This refinement facilitates the exploration of gene expression and regulation differences across developmental stages, breeds and tissues (Table 1). Furthermore, PIGOME offers nine practical tools designed to enhance the utilization of multi-omics datasets, a feature comparable to that of IAnimal (Table 2). Additionally, PIGOME facilitates the exploration of tissue-specific mRNA and ncRNA functional genomics in a more convenient manner, as we showed in the next section.

## Case study

Herein, we provide a case study to verify the usefulness of ‘Find tissue-specific genes’ in PIGOME and illustrate how to use PIGOME mining multi-omics information of interested genes. Initially, users can select the option ‘skeletal muscle’ in the ‘find tissue-specifically expressed gene’ section in the tool and thereafter, click the ‘Explore’ button. On the results page, based on substantial expression data, users can find 186 genes that are specifically expressed in skeletal muscle and then click the view icon of ‘*ENSSSCG00000026533’* to explore more detailed expression information about this gene (Figure 2A). On the expression page, users can obtain the gene symbol of myogenic factor 6 (*MYF6*), also known as *MRF4*, a myogenic regulatory factor involved in myogenesis. In addition, users can first see the related gene annotation and visualize its expression in different tissues using bar, line, or box-plot charts (Figure 2B and C). Importantly, users can also explore the expression trend of genes in skeletal muscles of different breeds and at different developmental stages (Figure 2D), demonstrating the ability to explore potential specific genes.

**Figure 2.**
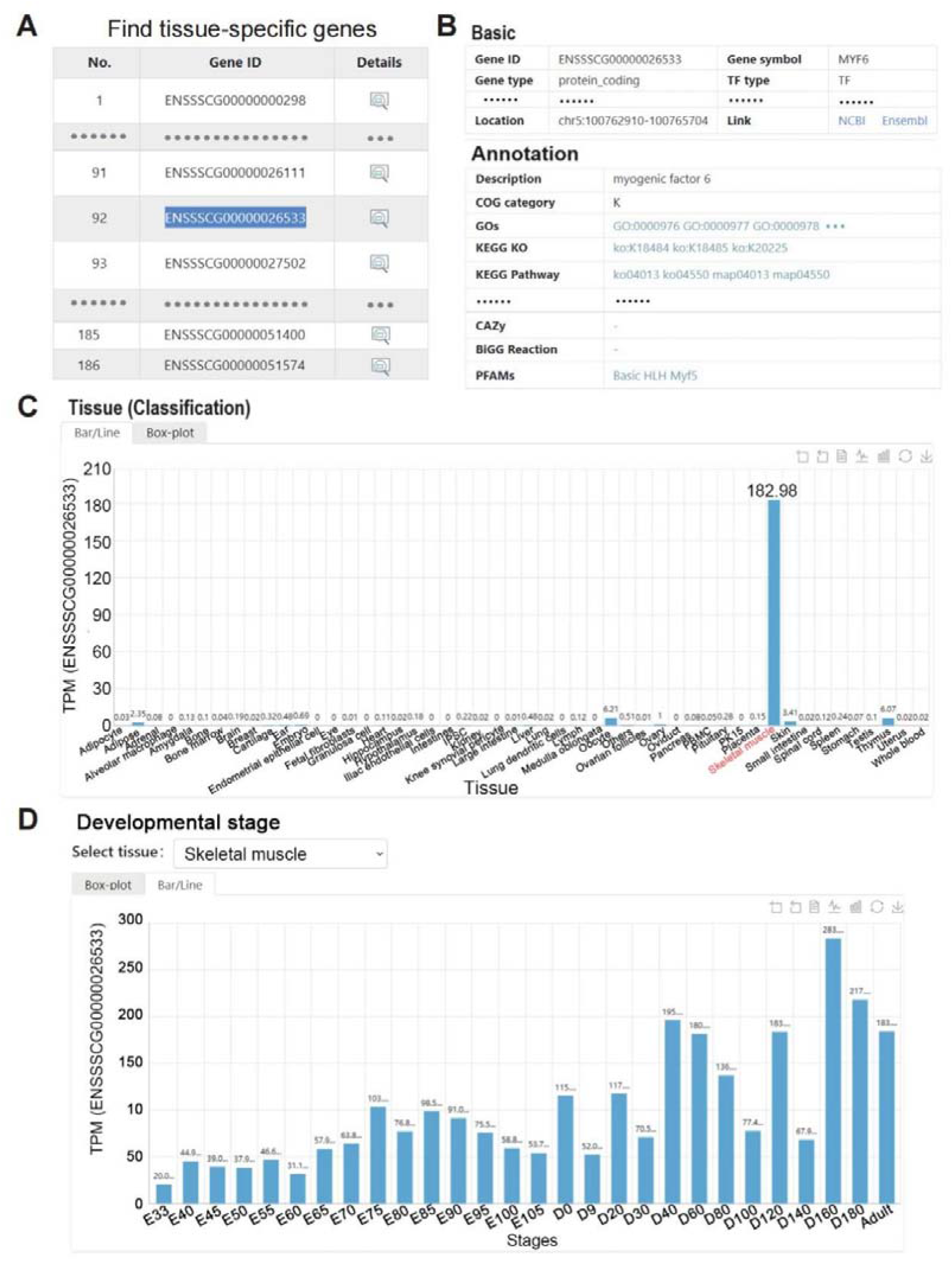
Tissue-specifically genes module in PIGOME. A. The results of finding tissue-specific genes in skeletal muscle. B. Annotation information related to *ENSSSCG00000026533* (*MYF6)*. C. The expression of *MYF6* in different tissues. D. Expression trend of *MYF6* in skeletal muscle at different developmental stages.

Finally, users can obtain all omics information related to genes using an exploration function. Users can use ‘*ENSSSCG00000026533*’ or ‘*MYF6*’ as the input in ‘Explore by gene ID or symbol’. The results page provides information, including SNP variation, expression abundance, annotation, epigenomics, and QTL related to *MYF6*. More importantly, one circRNA was identified in *MYF6* (Figure 3A). Serendipitously, by clicking the view icon (Figure 3B), it was shown that this circRNA (circ-MYF6), which may be a candidate circRNA that affects the development and growth of skeletal muscle, is also specifically expressed in skeletal muscle. There were 140 peaks identified from the ChIP-seq data, and 24 open chromatin regions were identified from ATAC-seq in *MYF6* (Figure 3C and D). Furthermore, the results showed 3,970 CpG methylation sites in exons, introns, and upstream regions (Figure 3E). Notably, we detected 24 SNPs and InDels in *MYF6* (Figure 3F). These results indicated that PIGOME can be used to explore the potential regulatory mechanisms of these genes.

**Figure 3.**
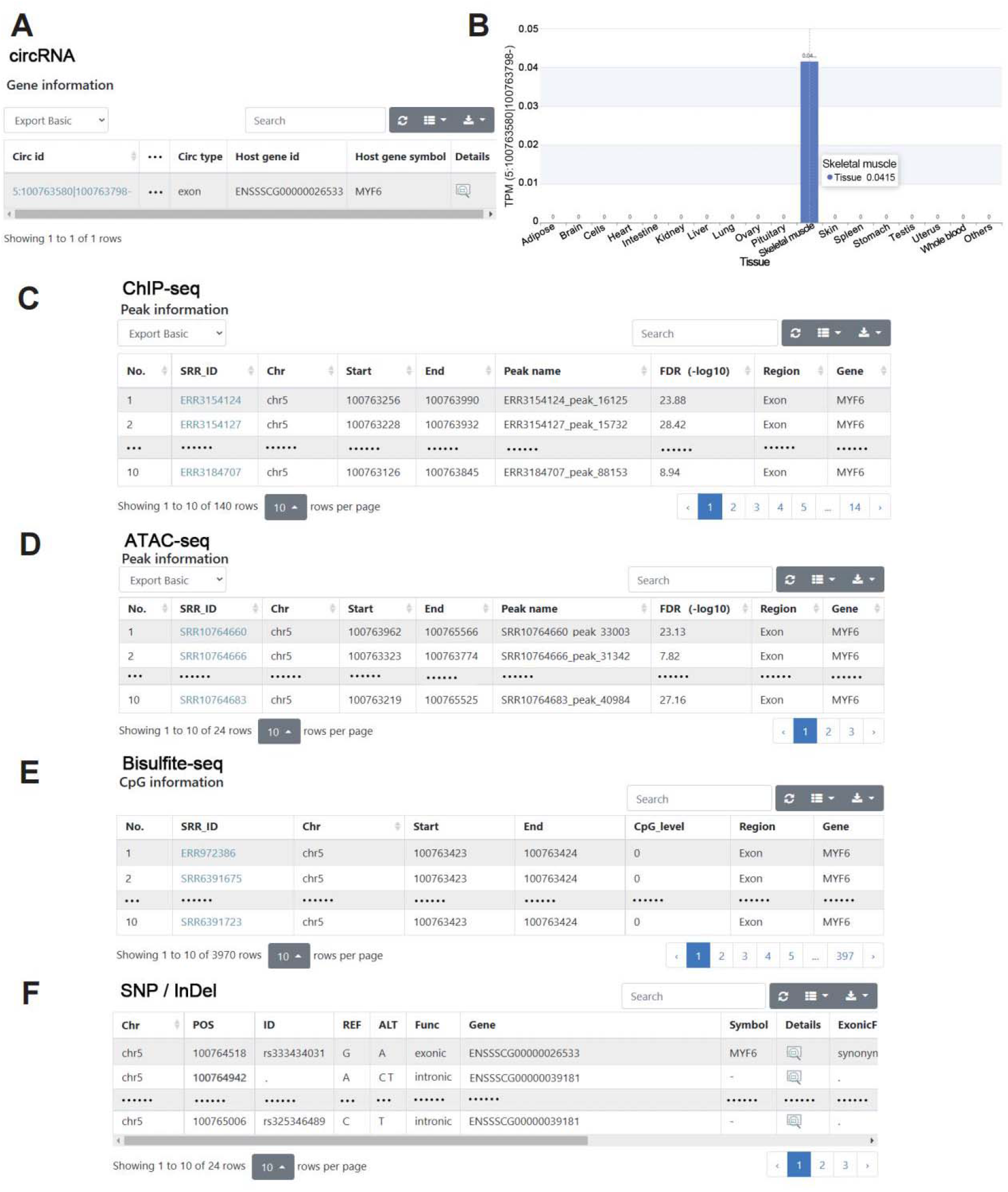
Explore the function and regulation of *MYF6* by PIGOME. A. The circRNA results are related to *MYF6*. B. Visualization results of circMYF6 expression in different tissues. C-F. The ChIP-seq (C), ATAC-seq (D), Bisulfite-seq (E), SNPs (F), and InDels (F) results related to *MYF6*, respectively.

## Discussion

In the past few decades, researchers have made great efforts in functional genomics research of pigs and accumulate valuable omics data [46]. Compared to earlier released databases for pigs, such as IAnimal [20], ISwine [21], and AOD [47], PIGOME contains the most data types and the most up-to-date multi-omics datasets covering comprehensive meta-information (Table 2). Moreover, PIGOME provides a user-friendly interface for browsing and analyzing omics data via interactive webpages, powerful search engines, and advanced tools. Integrated genomic, transcriptomic, and epigenomic data provide an efficient approach for discovering target genes and loci associated with economic traits and human-related diseases.

With the continuous development and innovation of high-throughput sequencing methods, more technologies have been developed, such as scRNA-seq and spatial transcriptomics [48,49]. The amount of omics data in public databases is also increasing. PIGOME will also constantly update new omics types and quantities. In the near future, other types of variations in the pig genome, such as structure variations (SVs), copy number variations (CNVs) and presence/absence variations (PAVs) will be available in PIGOME. We will aim to focus on cutting-edge single-cell sequencing and increase display related data, such as scRNA-seq, scATAC-seq and spatial transcriptome. In addition, PIGOME will update the latest gene annotation, genome-, epigenome- and transcriptome-wide association studies (GWAS, EWAS and TWAS) and applies quantitative trait locus (xQTL) information to help users better understand gene function. Furthermore, we will increase the internal relations among various data in the database and develop rich online tools. Finally, we intend PIGOME to be an important resource for exploring pig functional genomics, and will be of interest to the broad readership in the fields of animal genetics, breeding, and biomedical research.

## Data availability

PIGOME is available at https://pigome.com.

## CRediT author statement

**Guohao Han**: Methodology, Software, Visualization, Writing - Original Draft. **Peng Yang**: Methodology, Software, Data curation, Formal analysis. **Yongjin Zhang**: Data curation, Formal analysis. **Qiaowei Li**: Formal analysis. **Xinhao Fan**: Formal analysis. **Ruipu Chen**: Formal analysis. **Chao Yan**: Formal analysis. **Mu Zeng**: Formal analysis. **Yalan Yang**: Conceptualization, Project administration, Funding acquisition, Writing - review & editing. **Zhonglin Tang**: Conceptualization, Supervision, Funding acquisition, Writing - review & editing. All authors read and approved the final manuscript.

## Acknowledgments

We acknowledge the works of all the omics data researchers. This work was supported by the National Key Scientific Research Project [2023YFF1001100], the National Natural Science Foundation of China [U23A20229 and 32172697] and Agricultural Science and Technology Innovation Program [CAAS-ZDRW202006].

